## Supplementary Information for "mRNA trafficking directs cell-size-scaling of mitochondria distribution and function"

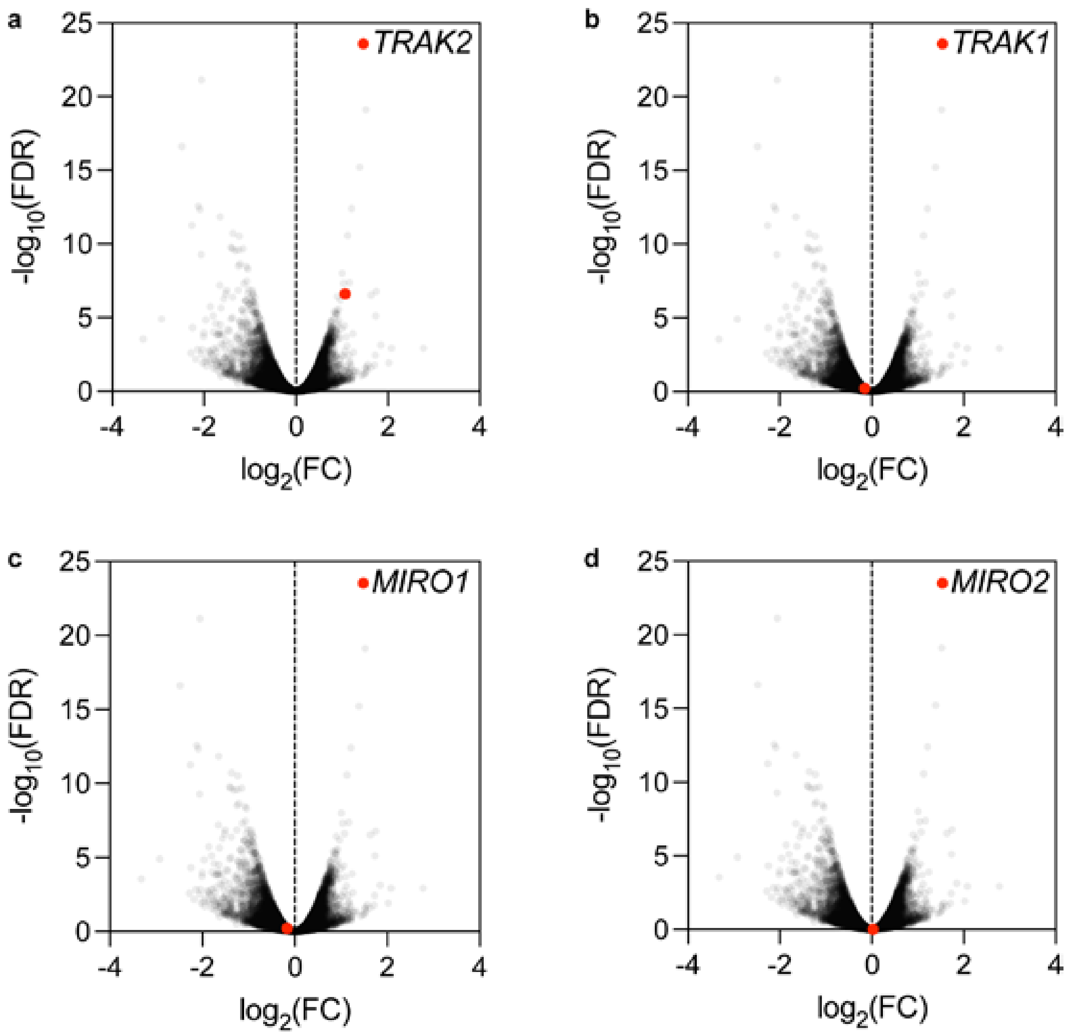
a
b
c
d
TRAK2
TRAK1
MIRO1
MIRO2
-log10(FDR)
log2(FC)
25
20
15
10
5
0
-4
-2
0
2
4

**Supplementary Fig. 1: *TRAK2* mRNA is polarised in migrating endothelial cells.**

**a**-**c**, Volcano plots of mRNAs differentially enriched in cell protrusions over cell bodies in migrating endothelial cells, as generated previously[1], with *TRAK2* (**a**), *TRAK1* (**b**), *MIRO1* (**c**) and *MIRO2* (**d**) indicated by red dots. RNAseq data are plotted in $\log_2$ fold change (FC) levels of protrusions over cell bodies against adjusted $-\log_{10}$ false discovery rate (FDR).

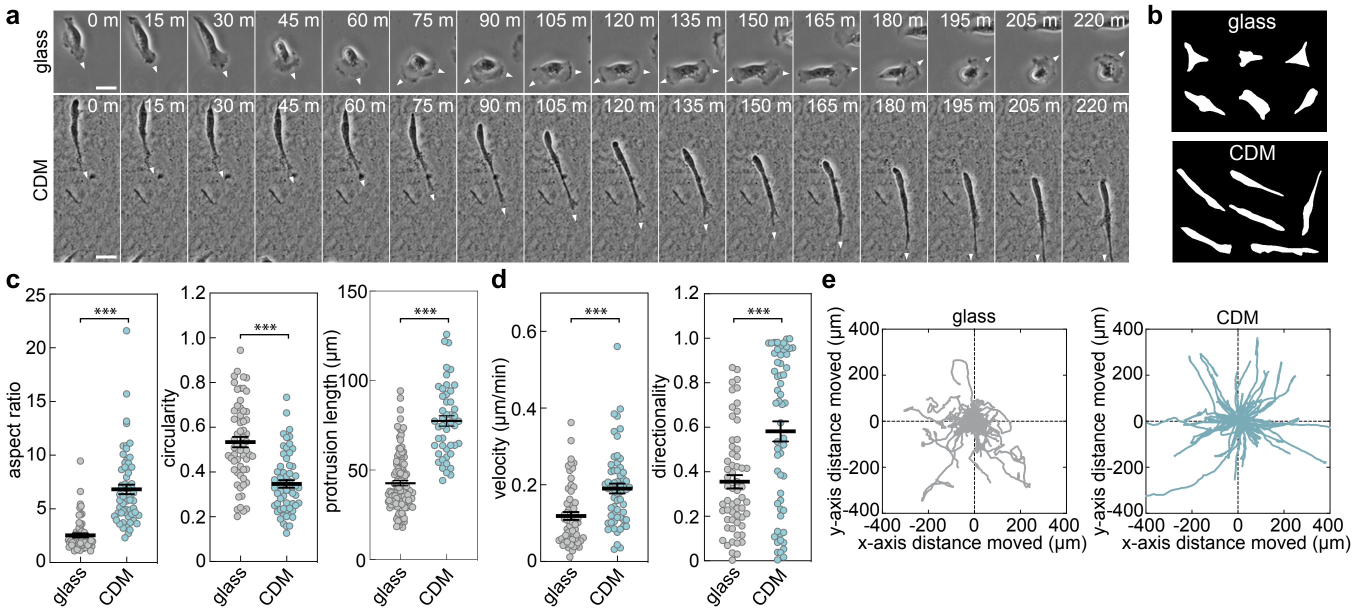

a
b
c
d
e
glass
CDM
0 m
15 m
30 m
45 m
60 m
75 m
90 m
105 m
120 m
135 m
150 m
165 m
180 m
195 m
205 m
220 m
aspect ratio
circularity
protrusion length (µm)
velocity (µm/min)
directionality
***
y-axis distance moved (µm)
x-axis distance moved (µm)

**Supplementary Fig. 2: Impact of substrate on cell morphology and behaviour.**

**a**, Still images of migrating human endothelial cells (ECs) cultured either on glass or CDM (arrowheads indicate direction of motile cell protrusions). **b**, Representative examples of EC morphology when cultured on glass or CDM. **c**, Quantification of EC morphometrics (aspect ratio, circularity and protrusion length) when cultured either on glass or CDM (*n*=60 at least cells glass, *n*=61 at least cells CDM, two-tailed Mann-Whitney test, ****P*=<0.0001). **d**, Quantification of EC velocity and directional persistence when cultured either on glass or CDM (*n*=60 cells glass, *n*=59 cells CDM, two-tailed Mann-Whitney test, ****P*=<0.0003). **e**, Rose plots of motile cell trajectories when cultured either on glass or CDM (*n*=60 cells glass, *n*=59 cells CDM). Scale bars, 20μm.

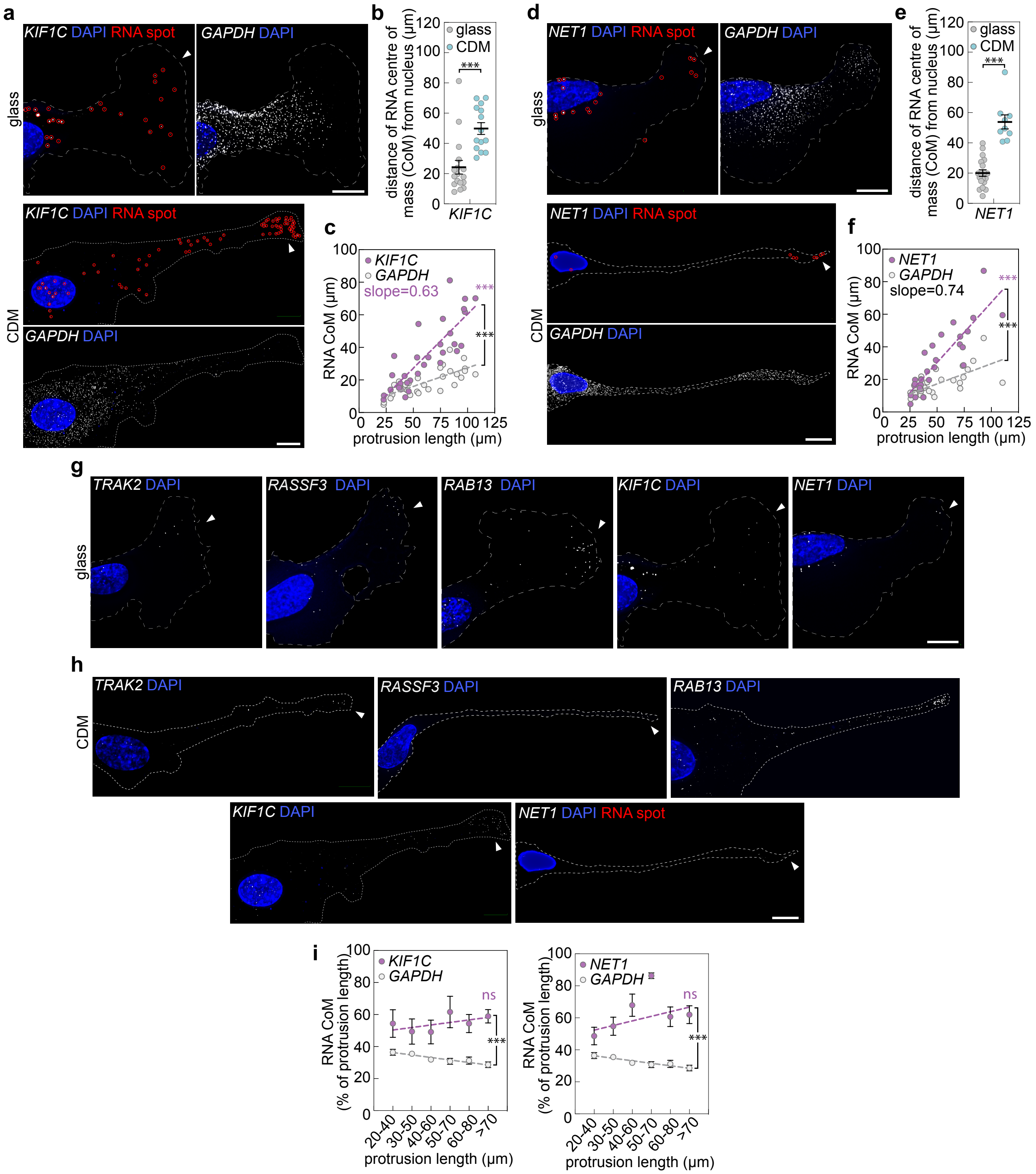

a
KIF1C DAPI RNA spot
GAPDH DAPI
glass
KIF1C DAPI RNA spot
CDM
GAPDH DAPI
b
distance of RNA centre of mass (CoM) from nucleus (μm)
glass
CDM
***
KIF1C
c
RNA CoM (μm)
KIF1C
GAPDH
slope=0.63
protrusion length (μm)
d
NET1 DAPI RNA spot
GAPDH DAPI
glass
NET1 DAPI RNA spot
CDM
GAPDH DAPI
e
distance of RNA centre of mass (CoM) from nucleus (μm)
glass
CDM
***
NET1
f
RNA CoM (μm)
NET1
GAPDH
slope=0.74
protrusion length (μm)
g
TRAK2 DAPI
RASSF3 DAPI
RAB13 DAPI
KIF1C DAPI
NET1 DAPI
glass
h
TRAK2 DAPI
RASSF3 DAPI
RAB13 DAPI
CDM
KIF1C DAPI
NET1 DAPI RNA spot
i
RNA CoM (% of protrusion length)
KIF1C
GAPDH
ns
***
protrusion length (μm)
20-40
30-50
40-60
50-70
60-80
>70
NET1
GAPDH

**Supplementary Fig. 3: Size-dependent targeting of *KIF1C* and *NET1* to cell protrusions.** **a**,**d**, smFISH detection of *KIF1C*, and *NET1* mRNAs in exemplar ECs cultured either on glass or CDM (red circles indicate distinct mRNA spots; arrowheads indicate mRNA accumulation at distal sites in protrusions; dashed line indicates cell outline). **b**,**e**, Quantification of the distance that the RNA centre of mass (CoM) sits from the nucleus for the indicated mRNAs when ECs were cultured either on glass or CDM (*n*=at least 17 cells glass, *n*=at least 9 cells CDM, two-tailed Mann-Whitney test, ****P*=<0.0001). **c**,**f**, Plots comparing the distance that the RNA CoM sits from the EC nucleus versus protrusion length for the indicated mRNAs (*n*=at least 28 cells; two-tailed Pearson's correlation coefficient, magenta asterisks, ****P*=<0.0001; analysis of covariance, black asterisks, ****P*=<0.0001 versus *GAPDH*). **g**,**h**, smFISH detection of *TRAK2, RASSF3, RAB13, KIF1C*, and *NET1* mRNAs in exemplar ECs (from Fig.1a,d,g and Supplementary Fig.3a,d) cultured either on glass (**g**) or CDM (**h**) without the labelling of RNA spots with red circles (arrowheads indicate mRNA accumulation at distal sites in protrusions; dashed line indicates cell outline). **i**, Plots comparing the distance that the RNA CoM sits from the EC nucleus normalised to protrusion length versus protrusion length for the indicated mRNAs (*n*=at least 28 cells; two-tailed Pearson's correlation coefficient, magenta asterisks, ns *P*=>0.2757; analysis of covariance, black asterisks, ****P*=<0.0038 versus *GAPDH*). For analyses in **c**,**f**,**i**, data for cells cultured on glass and CDM were pooled. Data are mean ± s.e.m. (**b**,**e**,**i**). For **a**,**d**,**g**,**h** scale bars, 10μm.

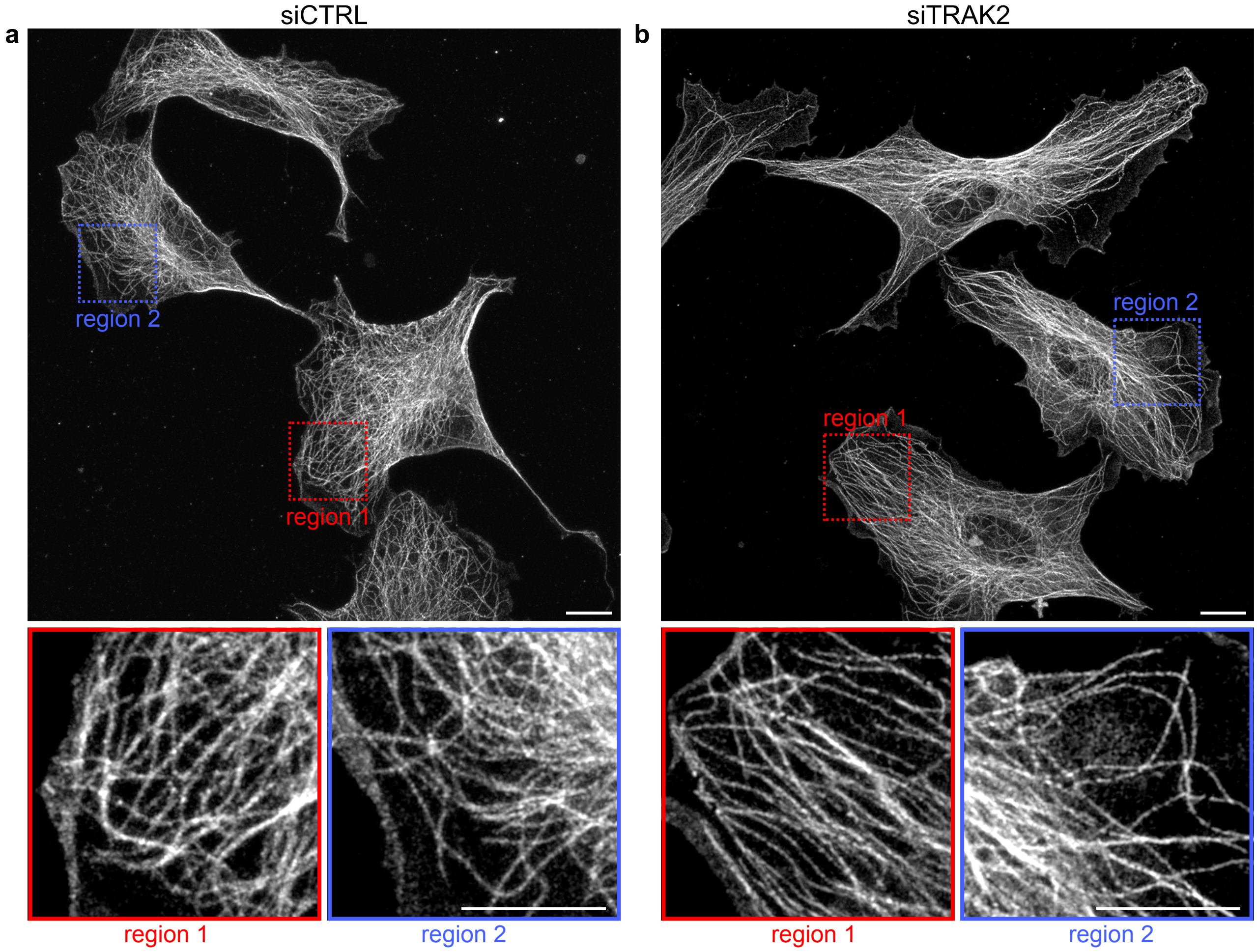
a
siCTRL
region 2
region 1
region 1
region 2
b
siTRAK2
region 2
region 1
region 1
region 2

**Supplementary Fig. 4: *TRAK2* knockdown does not perturb the microtubule cytoskeleton.** **a**,**b**, Representative images of alpha-tubulin-immunostained siCTRL (**a**) or siTRAK2 (**b**) treated ECs (red and blue boxes indicate the regions magnified below). Scale bars, 10μm.

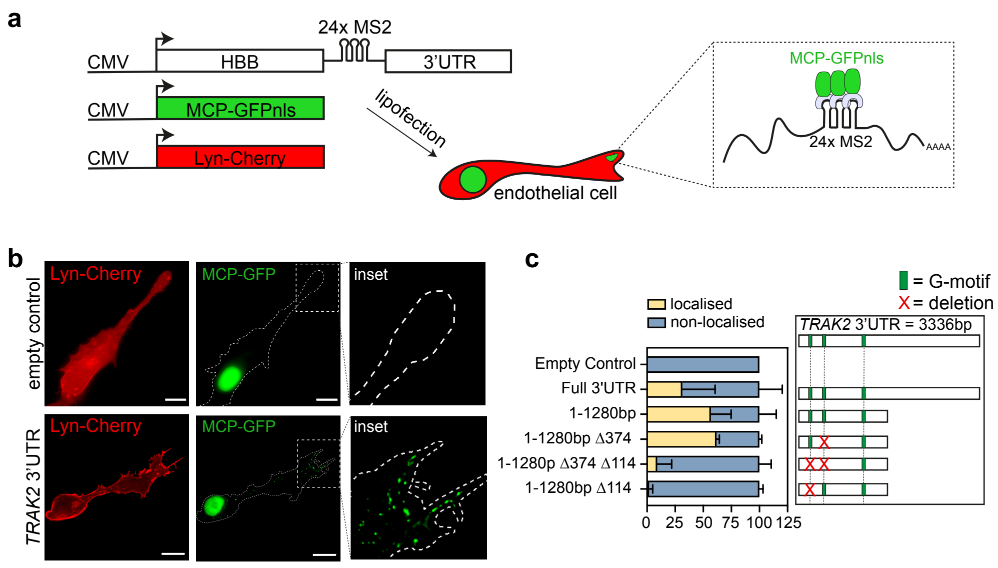
a
24x MS2
CMV
HBB
3'UTR
CMV
MCP-GFPnls
CMV
Lyn-Cherry
lipofection
endothelial cell
MCP-GFPnls
24x MS2
AAAA
b
empty control
Lyn-Cherry
MCP-GFP
inset
TRAK2 3'UTR
Lyn-Cherry
MCP-GFP
inset
c
= G-motif
= deletion
localised
non-localised
TRAK2 3'UTR = 3336bp
Empty Control
Full 3'UTR
1-1280bp
1-1280bp Δ374
1-1280p Δ374 Δ114
1-1280bp Δ114
0 25 50 75 100 125

**Supplementary Fig. 5: A single 3’UTR G-motif drives targeting of *TRAK2* mRNA.**

**a**, Schematic showing the *in vitro* MS2 system strategy used to identify 3’UTR elements driving mRNA localisation to protrusions. CMV promoter induced expression of lyn-Cherry, MCP-GFPnls, and hHBB-24xMS2-tagged 3′UTRs enables MCP-GFPnls to be used to visualise the localisation of 24xMS2-tagged 3’UTRs. **b**, Representative cells co-expressing lyn-Cherry, MCP-GFPnls, and either 24xMS2 or 24xMS2-*TRAK2* 3’UTR (dashed line indicates cell outline). **c**, Percentage of cells with MCP-GFPnls located in protrusions when transfected with either Wt *TRAK2* 3’UTR or the truncations and deletions of the *TRAK2* 3’UTR indicated in the panel to the right ($n$=3 independent experiments). This work is an extension of previous analyses[1]. Data are mean ± s.e.m. (**c**). Scale bars, 10μm.

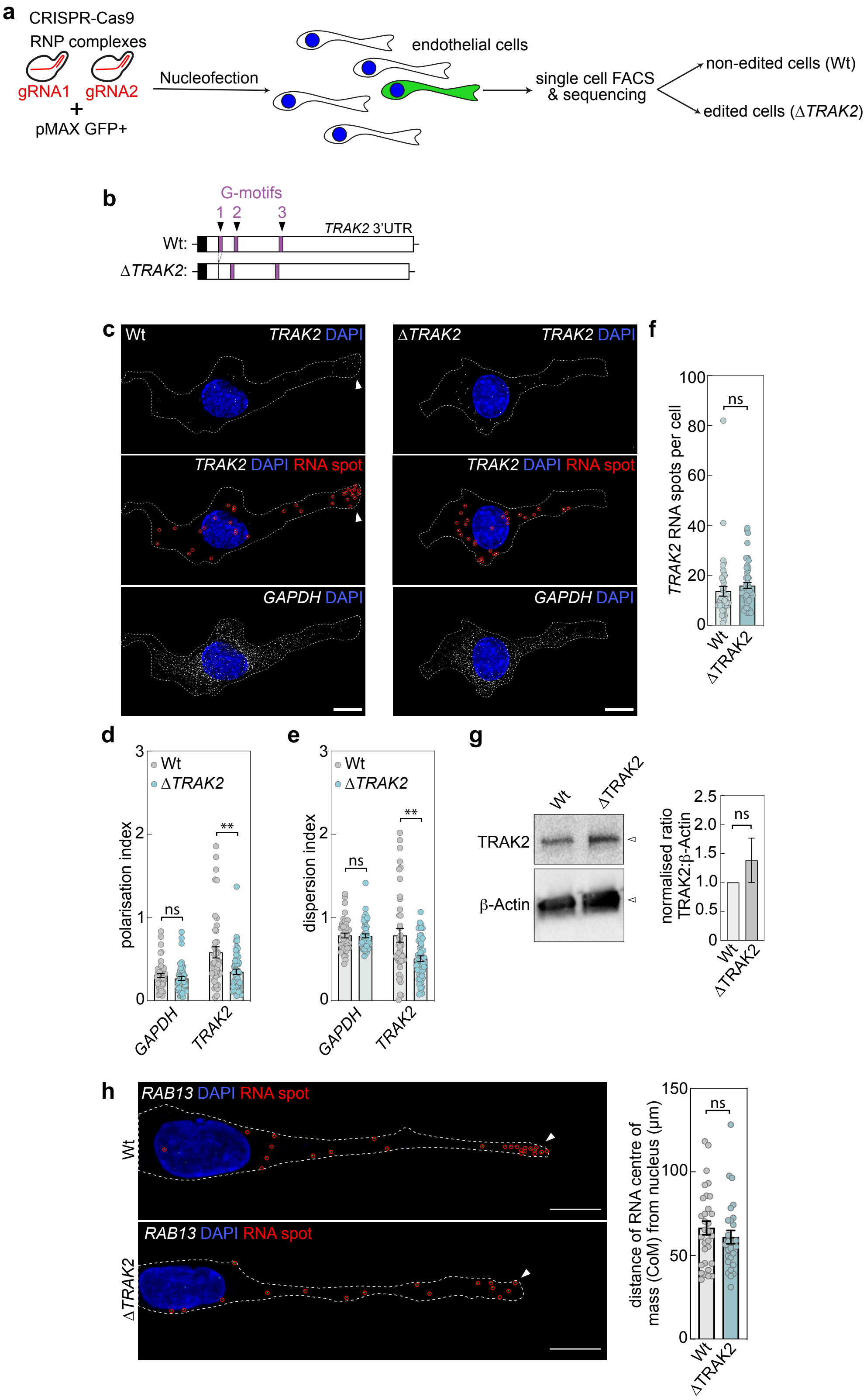
a
CRISPR-Cas9
RNP complexes
gRNA1
gRNA2
+
pMAX GFP+
Nucleofection
endothelial cells
single cell FACS
& sequencing
non-edited cells (Wt)
edited cells (ΔTRAK2)
b
G-motifs
1 2 3
TRAK2 3'UTR
Wt:
ΔTRAK2:
c
Wt
ΔTRAK2
TRAK2 DAPI
TRAK2 DAPI RNA spot
GAPDH DAPI
f
TRAK2 RNA spots per cell
ns
Wt
ΔTRAK2
d
polarisation index
Wt
ΔTRAK2
ns
**
GAPDH
TRAK2
e
dispersion index
ns
**
g
TRAK2
β-Actin
normalised ratio
TRAK2:β-Actin
h
RAB13 DAPI RNA spot
distance of RNA centre of
mass (CoM) from nucleus (μm)

**Supplementary Fig. 6: Deletion of the 24bp motif perturbs targeting of *TRAK2* mRNA.**

**a**, Schematic showing the strategy used to generate EC lines with both an excised 3'UTR localisation element (*ΔTRAK2*) and a non-edited control EC line (Wt). **b**, Illustration of removal of the 24bp G-motif from the endogenous *TRAK2* 3'UTR to generate *ΔTRAK2* (Black, white and magenta regions indicate the final exon of *TRAK2*, 3'UTRs and G-motifs, respectively). **c**, smFISH co-detection of *TRAK2* and *GAPDH* mRNAs in exemplar Wt or *ΔTRAK2* ECs cultured on glass (red circles indicate distinct mRNA spots; arrowheads indicate mRNA accumulation at distal sites in protrusions in Wt cells; dashed line indicates cell outline as defined by F-actin labelling). **d**,**e**, Quantification of the polarisation index (**d**) and dispersion index (**e**) of *TRAK2* and *GAPDH* mRNAs in Wt and *ΔTRAK2* ECs cultured on glass (*n*=44 cells Wt, *n*=at least 51 cells *ΔTRAK2*, two-tailed Mann-Whitney test, ***P*=<0.0096, ns *P*=>0.2373). **f**, Quantification of the number of *TRAK2* mRNA spots detected in Wt and *ΔTRAK2* ECs cultured on glass (*n*=44 cells Wt, *n*=52 cells *ΔTRAK2*, two-tailed unpaired t-test, ns *P*=0.3119). **g**, Representative Western blot for TRAK2 and β-Actin in Wt and *ΔTRAK2* ECs and densitometric analysis the ratio of TRAK2:β-Actin levels (*n*=4, two-tailed Mann-Whitney test, ns *P*=0.3143). **h**, smFISH detection of *RAB13* mRNAs in exemplar Wt or *ΔTRAK2* ECs cultured on CDM (red circles indicate distinct mRNA spots; arrowheads indicate mRNA accumulation at distal sites in protrusions; dashed line indicates cell outline as defined by F-actin labelling) and quantification of the RNA CoM (*n*=31 cells Wt, *n*=28 cells *ΔTRAK2*, two-tailed Mann-Whitney test, ns *P*=0.3282). Data are mean ± s.e.m. (**d**,**e**,**f**,**g**,**h**). For **c** and **h** scale bars, 10μm.

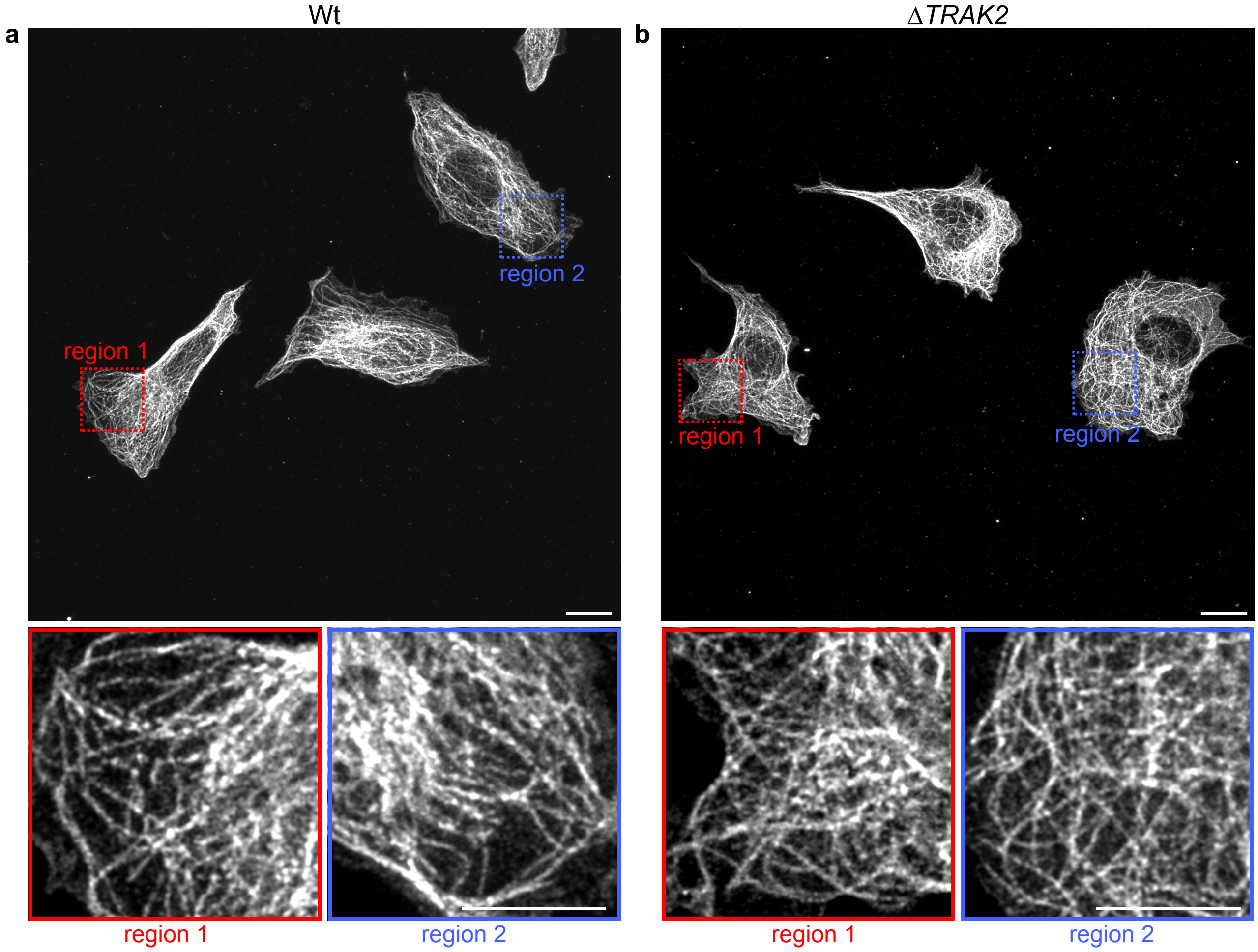

a
Wt
region 1
region 2
b
ΔTRAK2
region 1
region 2
region 1
region 2
region 1
region 2

**Supplementary Fig. 7: Loss*TRAK2* mRNA targeting does not perturb the microtubule cytoskeleton. a**,**b**, Representative images of alpha-tubulin-immunostained Wt (**a**) or *ΔTRAK2* (**b**) ECs (red and blue boxes indicate the regions magnified below). Scale bars, 10μm.

**Supplementary table 1: Oligonucleotide Sequences.**

| | |
|---|---|
| **CRISPr/Cas9** | |
| **Name** | **Sequences (5' – 3')** |
| 5' crRNA *TRAK2* 3' UTR | AGAAAAGGAATGTTGCACAA |
| 3' crRNA *TRAK2* 3' UTR | GGAGGAATGGAAATAAAATT |
| **Genotyping** | |
| **Name** | **Sequences (5' – 3')** |
| *TRAK2* 3' UTR F | TGAAACATGTGGTCTGGTCTG |
| *TRAK2* 3' UTR R | TCCCTCTGAACACTCATGGC |
| **Cloning** | |
| **Name** | **Sequences (5' – 3')** |
| NheI 1nt *TRAK2* 3'UTR F | GGAAGCTAGCGGTTCAGCAGTTAACTGACC |
| XhoI 3336nt *TRAK2* 3'UTR R | GGAACTCGAGTGAAAATTCAAGCTACTCATG |
| XhoI 1280nt *TRAK2* 3'UTR R | GGAACTCGAGATCTAACCCCAGAGCCTCAC |
| del114-142 *TRAK2* R | GGGCTCATCCCAATTTTATCACAACCCTTGTGCAACAT |
| del114-142 *TRAK2 F* | ATGTTGCACAAGGGTTGTGATAAAATTGGGATGAGCCC |
| del374-402 *TRAK2* F | AAGCATAAAGCAGAGAGAACCCCAGTTTTATTGCTTATAGAAAGC |
| del374-402 *TRAK2* R | GCTTTCTATAAGCAATAAAACTGGGGTTCTCTCTGCTTTATGCTT |
| **Notes:** underlined uppercase - restriction site | |

### Supplementary table 2: Probe Sequences.

| mRNA | Probe sequence (5' - 3') | | mRNA | Probe sequence (5' - 3') | | mRNA | Probe sequence (5' - 3') |
|---|---|---|---|---|---|---|---|
| *ACTB* | gctcgagccataaaaggcaa | | *KIF1C* | tcttctatcacctactcgg | | *NET1* | gccgagtcagaaccaaga |
| *ACTB* | cgatatcatcatccatggtg | | *KIF1C* | ttggtttgctcctggcaac | | *NET1* | ccgttcgttccgtgtgac |
| *ACTB* | cacgatggaggggaagacgg | | *KIF1C* | cttcctatctctcttcttca | | *NET1* | gccggtaaacctggtaag |
| *ACTB* | acataggaatccttctgacc | | *KIF1C* | cagcgtttgtgggacacac | | *NET1* | ctcttggactgggattgg |
| *ACTB* | ggtacttcagggtgaggatg | | *KIF1C* | gtcaaacactggctacgca | | *NET1* | tgcaggtcttctaggact |
| *ACTB* | cagattttctccatgtcgtc | | *KIF1C* | acttatgatcctaccatgc | | *NET1* | ccattctcacatctccat |
| *ACTB* | acacgcagctcattgtagaa | | *KIF1C* | ccaaatatagcaggaggcc | | *NET1* | agctcctcgaaaggagcc |
| *ACTB* | acatgatctgggtcatcttc | | *KIF1C* | cagccaaaccagtcaggtc | | *NET1* | gctttctctgagttactg |
| *ACTB* | ggatagcacagcctggatag | | *KIF1C* | tattaaagggcccacatcc | | *NET1* | gaggggtcatggaagcga |
| *ACTB* | catcacgatgccagtggtac | | *KIF1C* | aagctccctgtttgctatc | | *NET1* | gagtgtgagactgggctg |
| *ACTB* | tcgtagatgggcacagtgtg | | *KIF1C* | aaagatcgggctgcgcacc | | *NET1* | gtggaacacgtcattggc |
| *ACTB* | tcttcatgaggtagtcagtc | | *KIF1C* | cttctgtgctttgtccata | | *NET1* | cagttgaaccactgctgc |
| *ACTB* | taatgtcacgcacgatttcc | | *KIF1C* | aagcattgacaggctggcg | | *NET1* | cgactggaagggggcaat |
| *ACTB* | atctcttgctcgaagtccag | | *KIF1C* | atctgtcagcaacagccag | | *NET1* | tgcagctcaggtggactg |
| *ACTB* | cagggaggagctggaagcag | | *KIF1C* | gtaccccacaaatccttct | | *NET1* | tcacactcttcgtgcagc |
| *ACTB* | tcattgccaatggtgatgac | | *KIF1C* | gacttaaagcagaactcct | | *NET1* | tgagtttcctcgcagagg |
| *ACTB* | gaaggtagtttcgtggatgc | | *KIF1C* | gcttgggagacctttcaag | | *NET1* | gaaactgtggatgccctc |
| *ACTB* | cgtcacacttcatgatggag | | *KIF1C* | aggtaagcgcagtatgctt | | *NET1* | gcgttttcatcaacttct |
| *ACTB* | tacaggtctttgcggatgtc | | *KIF1C* | gcaggtaacacaaccagga | | *NET1* | atgccagagccacatctg |
| *ACTB* | caatgccagggtacatggtg | | *KIF1C* | aacaatgatgtagcctgcc | | *NET1* | agctcttgctgtcctctg |
| *ACTB* | atcttcattgtgctgggtgc | | *KIF1C* | gcctcaagccagaacatta | | *NET1* | ctgtgtctggtgtgtctt |
| *ACTB* | ctcaggaggagcaatgatct | | *KIF1C* | tggcatagtcaagggctcg | | *NET1* | tccctcgctcttcggatg |
| *ACTB* | cgatccacacggagtacttg | | *KIF1C* | acctgcaaaatcctggcaa | | *NET1* | tgccaccagaaagggctt |
| *ACTB* | tcatactcctgcttgctgat | | *KIF1C* | gcagtgctgtagtacaatc | | *NET1* | caccaaagtctctttccg |
| *ACTB* | atttgcggtggacgatggag | | *KIF1C* | ccagagtctggcaagccaa | | *NET1* | aacacacagagccttctc |
| *ACTB* | aagtcatagtccgcctagaa | | *KIF1C* | aagcagtagctcagagctt | | *NET1* | gtaaccctatgatgctcc |
| *ACTB* | gtcaagaaagggtgtaacgc | | *KIF1C* | ctatggttttggctggaga | | *NET1* | taccaggcttaccatgtt |
| *ACTB* | ttttctgcgcaagttaggtt | | *KIF1C* | aacctaatcgttccttcgc | | *NET1* | agccatgatgaatccaga |
| *ACTB* | cattgtgaactttgggggat | | *KIF1C* | aaaggcaggagcatcaggg | | *NET1* | actggcagaattccagca |
| *ACTB* | gtgcaatcaaagtcctcggc | | *KIF1C* | ggcagtcagaggtgtcaaa | | *NET1* | aggttttgggtgccaatg |
| *ACTB* | cctgtaacaacgcatctcat | | *KIF1C* | ctgtcaacaggtctgtgaa | | *NET1* | ctccagacatacattcca |
| *ACTB* | cttttaggatggcaagggac | | *KIF1C* | tacattccagatcttgagt | | *NET1* | tggtttgttttccaagga |
| *ACTB* | tctccttagagagaagtggg | | *KIF1C* | cactgttgccaaaccatcc | | *NET1* | taaaaacctcctcaggct |
| *ACTB* | gtggacttgggagaggactg | | *KIF1C* | tctatcattttcttgctca | | *NET1* | ggcttggtttcacttggt |
| *ACTB* | aaagcaatgctatcacctcc | | *KIF1C* | gcattctcggtaacactca | | *NET1* | ccctgaaatgctacactt |
| *ACTB* | catacatctcaagttggggg | | *KIF1C* | ttttctcatctcaaacccc | | *NET1* | agccatttcatgggtctt |
| *ACTB* | actcccagggagaccaaaag | | *KIF1C* | tctaagtaaagtttgccca | | *NET1* | ccctcttgcatttcagac |
| *ACTB* | gtctcaagtcagtgtacagg | | *KIF1C* | cttgatggcctagaggggt | | *NET1* | cccccttttcaaagatgat |
| *ACTB* | ggtgtgcacttttattcaac | | *KIF1C* | gaggtgatggtgatgtctg | | *NET1* | atcgggcttttactcctc |
| | | | *KIF1C* | aaggccccatggattctaa | | *NET1* | ttatcatttccactggcc |

| mRNA | Probe sequence (5' - 3') | | mRNA | Probe sequence (5' - 3') | | mRNA | Probe sequence (5' - 3') |
|---|---|---|---|---|---|---|---|
| *RAB13* | caggccaggaagaagttttc | | *RASSF3* | ggtgaagaagaagtcctcgg | | *TRAK2* | cacacaattggggaagccaa |
| *RAB13* | tggacggttggcaaacagag | | *RASSF3* | ttcaacttgtctgtgactgc | | *TRAK2* | ctttggacagtcatgactgg |
| *RAB13* | cagcaacttgaagaggtggt | | *RASSF3* | cccatttgaattcaaggtca | | *TRAK2* | gcactctcatgtgattatca |
| *RAB13* | atgatcagacaagtcttgcc | | *RASSF3* | cagagttccatctgtacttt | | *TRAK2* | gtcaaatgacatccgtagca |
| *RAB13* | gttgttgaagttgtcctctg | | *RASSF3* | tggagaagtctgtggaggtt | | *TRAK2* | aaatggatttggtcacatgg |
| *RAB13* | ttccgatggtggagatgtaa | | *RASSF3* | tgggagagagttttccagaa | | *TRAK2* | ggaaatccatcaagccattc |
| *RAB13* | ccacagtgcggatcttgaaa | | *RASSF3* | tgtattcatacagccattgc | | *TRAK2* | gctttggtatgaatcagagt |
| *RAB13* | ccagacttgtagtttgatct | | *RASSF3* | tgtttgtgctgctgatatga | | *TRAK2* | ctttcttgctttggttgaga |
| *RAB13* | attgtcttgaaccgctcttg | | *RASSF3* | gctctcagtcacgagaaact | | *TRAK2* | attctgggattgactcatgc |
| *RAB13* | tccacggtagtaggcagtag | | *RASSF3* | ccctgtgacaacgcttataa | | *TRAK2* | ttcacctgttggtgatgtaa |
| *RAB13* | atactaggataatgcccatg | | *RASSF3* | taccaaacgcaggtagagtg | | *TRAK2* | gagtctctgtgattgctatt |
| *RAB13* | gatttctcatccgtgatgtc | | *RASSF3* | actaagtgtgtctgttctgg | | *TRAK2* | cattggagcagacatcagtg |
| *RAB13* | ccagttctgaatattctcga | | *RASSF3* | ctccaatttcatgttcacga | | *TRAK2* | agctcaacttcagggagatc |
| *RAB13* | cattctccttgatgcttttc | | *RASSF3* | tagttctggaaggctgaagg | | *TRAK2* | gtagttgttcttctagcaga |
| *RAB13* | ctccatgtcacatttgttcc | | *RASSF3* | ttgtccaagatgcgcaagaa | | *TRAK2* | agtgtctacttttagcctat |
| *RAB13* | aaatcggattccatgctctc | | *RASSF3* | ctttaatcaggcttccacac | | *TRAK2* | agtccagtcttgattttcat |
| *RAB13* | ctggatttagcactagtttc | | *RASSF3* | ttgctgctgttttgtacatc | | *TRAK2* | aacggaaagtctcttcagca |
| *RAB13* | aaaagcctcatccacattca | | *RASSF3* | gatagaaaggcgaagaccct | | *TRAK2* | ctgtctgtgcctagaatcat |
| *RAB13* | ctcctgacttgagcaagatg | | *RASSF3* | gaccaggtttggggaacaaa | | *TRAK2* | gtaagttttggtcatctgct |
| *RAB13* | agttttcaggtcagtactgg | | *RASSF3* | ttacatgggacgctaaggtg | | *TRAK2* | ggagatgtgtaaccatgtcg |
| *RAB13* | acttgttggtgttcttcttg | | *RASSF3* | gtgctgaacaacaagcttgc | | *TRAK2* | cagatcacgatccctctctg |
| *RAB13* | aggcaagaaagggtcctcag | | *RASSF3* | cttgccaagaggtaacactg | | *TRAK2* | tgtccaattcgagcagcgag |
| *RAB13* | tacctatgtgaccctccaag | | *RASSF3* | cctttcgattcacatggata | | *TRAK2* | acatggttccgctttaagag |
| *RAB13* | aacccaggtaaggtctgaag | | *RASSF3* | aattcctcagagttcctaga | | *TRAK2* | ggattcgttctgctcagata |
| *RAB13* | tttacatttatgtttgccct | | *RASSF3* | actgaaaagccaagtcacct | | *TRAK2* | tcaaaggcttgtcccaattg |
| *RAB13* | ggaccctaaaacctgatcta | | *RASSF3* | aatgggtttaacacctccac | | *TRAK2* | atgctgcagctgattaactt |
| *RAB13* | gagcaaattccctagtgtag | | *RASSF3* | atattctacccaccaggaaa | | *TRAK2* | actcatctttcttgcatagc |
| *RAB13* | agaccatgacaagtgacaga | | *RASSF3* | gggcagaattaaagggtcgg | | *TRAK2* | cttcagaagcaatggagacg |
| *RAB13* | tgcaaatggtggcctttaat | | *RASSF3* | aatagccctgttctctatta | | *TRAK2* | cagctggaatcagtttcact |
| | | | *RASSF3* | tggtaggattgggtctttat | | *TRAK2* | tcattgaaccgaagaggtgt |
| | | | *RASSF3* | ttctattgcacatgcatctc | | *TRAK2* | gcaacccttgagataagcta |
| | | | *RASSF3* | gaggttatttagcctgtttt | | *TRAK2* | ttttcttgcagcatttccaa |
| | | | *RASSF3* | ccctctgttctcagattaaa | | *TRAK2* | catattctcttcttccagtt |
| | | | *RASSF3* | aatccattaggtgtcaccaa | | *TRAK2* | gacaagccttggatcgaaga |
| | | | *RASSF3* | ctgaagcaccttggaatttc | | *TRAK2* | aggtaacagtttctgtcttt |
| | | | *RASSF3* | aaaggcacagttcacttggg | | *TRAK2* | agctgttgttccttttcttc |
| | | | *RASSF3* | cagaagagccaccaagagac | | *TRAK2* | ttctttaacacagtcgctga |
| | | | *RASSF3* | ttaagtctgcctccaacaag | | *TRAK2* | tctgagcatttgtttcacga |
| | | | *RASSF3* | gtaaattgtgtggcacagca | | *TRAK2* | gaatcagctcatcactcttc |
| | | | *RASSF3* | gaaggcattccctgaacaat | | *TRAK2* | agaggaaagctcttcttggt |
| | | | *RASSF3* | atactcaaatccaggccata | | *TRAK2* | aggtctacaatctgtgacaa |
| | | | *RASSF3* | ccatcagctactactttagg | | *TRAK2* | atgttctttaagtttgtgct |
| | | | *RASSF3* | gatgaggatgcctattcaga | | *TRAK2* | catctttggaagcttgcagg |
| | | | *RASSF3* | cacatgacagtaagcgaggt | | *TRAK2* | ctgtcttgtaactcgtgcag |
| | | | *RASSF3* | accagtgtttgggaaaggac | | *TRAK2* | aacattcctagacactccat |
| | | | *RASSF3* | aataccacccccttcatttaa | | *TRAK2* | gggccagatctactacgaag |
| | | | *RASSF3* | tgccatttaaaatgctctcc | | *TRAK2* | cccagtaaaagctccatatg |
| | | | | | | *TRAK2* | caatctcagctgccaaagat |
